# *Enterococcus faecalis* CRISPR-Cas is a robust barrier to conjugative antibiotic resistance dissemination in the murine intestine

**DOI:** 10.1101/312751

**Authors:** Valerie J. Price, Sara W. McBride, Karthik Hullahalli, Anushila Chatterjee, Breck A. Duerkop, Kelli L. Palmer

## Abstract

CRISPR-Cas systems are barriers to horizontal gene transfer (HGT) in bacteria. Little is known about CRISPR-Cas interactions with conjugative plasmids, and studies investigating CRISPR-Cas/plasmid interactions in *in vivo* models relevant to infectious disease are lacking. These are significant gaps in knowledge because conjugative plasmids disseminate antibiotic resistance genes among pathogens *in vivo*, and it is essential to identify strategies to reduce the spread of these elements. We use enterococci as models to understand the interactions of CRISPR-Cas with conjugative plasmids. *Enterococcus faecalis* is a native colonizer of the mammalian intestine and harbors pheromone-responsive plasmids (PRPs). PRPs mediate inter- and intraspecies transfer of antibiotic resistance genes. We assessed *E. faecalis* CRISPR-Cas anti-PRP activity in the mouse intestine and under varying *in vitro* conditions. We observed striking differences in CRISPR-Cas efficiency *in vitro* versus *in vivo*. With few exceptions, CRISPR-Cas blocked intestinal PRP dissemination, while *in vitro*, the PRP frequently escaped CRISPR-Cas defense. Our results further the understanding of CRISPR-Cas biology by demonstrating that standard *in vitro* experiments do not adequately model the *in vivo* anti-plasmid activity of CRISPR-Cas. Additionally, our work identifies several variables that impact the apparent *in vitro* anti-plasmid activity of CRISPR-Cas, including planktonic versus biofilm settings, different donor/recipient ratios, production of a plasmid-encoded bacteriocin, and the time point at which matings are sampled. Our results are clinically significant because they demonstrate that barriers to HGT encoded by normal human microbiota can have significant impacts on *in vivo* antibiotic resistance dissemination.

**Importance:** CRISPR-Cas is a type of immune system encoded by bacteria that is hypothesized to be a natural impediment to the spread of antibiotic resistance genes. In this study, we directly assessed the impact of CRISPR-Cas on antibiotic resistance dissemination in the mammalian intestine and under varying *in vitro* conditions. We observed a robust effect of CRISPR-Cas on *in vivo* but not *in vitro* dissemination of antibiotic resistance plasmids in the native mammalian intestinal colonizer *Enterococcus faecalis*. We conclude that standard laboratory experiments currently do not appropriately model the *in vivo* conditions where antibiotic resistance dissemination occurs between *E. faecalis* strains. Moreover, our results demonstrate that CRISPR-Cas encoded by native members of the mammalian intestinal microbiota can block the spread of antibiotic resistance plasmids.

## Introduction

CRISPR-Cas systems confer adaptive immunity against mobile genetic elements (MGEs) in bacteria (1–3). CRISPR-Cas systems utilize nucleases programmed with small RNAs to direct sequence-specific cleavage of nucleic acids including phage and plasmids (4). Most experimental studies of native CRISPR-Cas systems have examined either anti-phage defense or defense against electrotransformed plasmids in low complexity *in vitro* systems. Comparatively little information is available on the roles of CRISPR-Cas in regulating plasmid conjugation, and there have been few experimental studies assessing the function of CRISPR-Cas systems within the native ecology of microbial communities. These are major weaknesses in the field from a public health perspective. Conjugative plasmids disseminate antibiotic resistance genes, and CRISPR-Cas systems are naturally occurring barriers that could impede the dissemination of these genes in mammalian microbiota.

We use *Enterococcus faecalis* as a model organism to study the interactions of CRISPR-Cas systems with conjugative plasmids. *E. faecalis* is a gram-positive bacterium, a native inhabitant of the mammalian intestine (5), and an opportunistic pathogen that is among the leading causes of hospital-acquired infections (HAIs) in the United States (6, 7). *E. faecalis* strains causing HAIs possess unique characteristics relative to strains that normally colonize the human intestine. HAI strains typically have larger genomes resulting from rampant plasmid, phage, and other MGE acquisition (8, 9). Multidrug-resistant (MDR) *E. faecalis* generally lack CRISPR-Cas systems, and there is a correlation between the absence of CRISPR-Cas and the presence of horizontally acquired antibiotic resistance in *E. faecalis* clinical isolates (10). From genomic analyses, it appears that CRISPR-Cas is a potent barrier to the horizontal acquisition of antibiotic resistance in *E. faecalis*. Our subsequent efforts have attempted to experimentally address this hypothesis.

The model plasmids we use for our studies are the pheromone-responsive plasmids (PRPs). The PRPs appear to be highly co-evolved with *E. faecalis* (11, 12). PRPs are large (can be >60 kb) and encode accessory traits such as antibiotic resistance, bacteriocin production, reduced UV light susceptibility, and enhanced biofilm formation (11). PRPs encoding antibiotic resistance genes are often present in *E. faecalis* infection isolates (11, 13–15). The model PRP, pAD1, encodes genes for production and self-immunity to a bacteriocin called cytolysin (16). Cytolysin is a lantibiotic-like antimicrobial peptide and hemolysin with activity against a number of gram-positive bacteria (17, 18).

In this study, we utilize *E. faecalis* T11RF, a non-MDR strain that encodes a Type II CRISPR-Cas system referred to as CRISPR3-Cas (10, 19). Type II CRISPR-Cas systems employ a Cas9-crRNA-tracrRNA ribonucleoprotein complex to generate double-stranded DNA breaks in invading MGEs (3, 20, 21). Sequence specificity in the cleavage event is conferred by the crRNA (22). A crRNA is encoded by a short sequence referred to as a spacer, which is derived from and is complementary to a previously encountered MGE (1, 23, 24). The *E. faecalis* T11RF CRISPR3-Cas system encodes a spacer with perfect sequence complementarity to the *repB* gene of the PRP pAD1 (10, 19). In previous studies, we demonstrated that the *E. faecalis* CRISPR3-Cas system interferes with the conjugative acquisition of pAM714 (19), a pAD1 variant with an insertion of Tn*917* encoding *ermB* (25, 26). More specifically, pAM714 acquisition is decreased by ∼80-fold in *E. faecalis* T11RF relative to T11RFΔ*cas9* after 18 hours biofilm mating on an agar surface, and CRISPR3-Cas defense against pAM714 requires the targeting spacer (19). These results support our overarching hypothesis that CRISPR-Cas is a significant barrier to the horizontal acquisition of antibiotic resistance in *E. faecalis*. However, the magnitude of CRISPR3-Cas impact on pAM714 acquisition, while significant, was low compared with the overall high transfer rate of pAM714 under the conditions tested. Many pAM714 molecules escaped CRISPR3-Cas defense despite T11RF possessing functional CRISPR-Cas. We have made similar observations in other *E. faecalis* strains using both native and engineered CRISPR-Cas systems and with both naturally occurring and engineered plasmids (19, 27, 28). We previously defined the ability of cells to acquire CRISPR-targeted plasmids at high frequencies as CRISPR tolerance (27).

To investigate potential explanations for the seeming discrepancy between the presence of CRISPR-Cas in wild *E. faecalis* isolates and our *in vitro* observations of statistically significant but middling population-level impact of CRISPR-Cas on conjugative plasmid transfer, we first assessed whether different *in vitro* mating conditions alter conclusions reached about CRISPR-Cas defense efficiency. We compared pAM714 acquisition by wild type and Δ*cas9* T11RF recipients in planktonic and agar plate biofilm matings using time course experiments and two different initial donor/recipient ratios. We performed the same experiments with the PRP pAM771, which is a pAD1 derivative possessing a Tn*917* insertion in the cytolysin locus (25, 29, 30). We reasoned that killing of plasmid-free recipient cells by the cytolysin could ‘punish’ cells that utilize CRISPR-Cas against the plasmid, potentially altering the apparent efficacy of CRISPR-Cas. We also assessed the transfer of pAM714 and pAM771 to wild-type and Δ*cas9* T11RF recipients in a murine intestinal colonization model. To our knowledge, this is the first study to assess the impact of CRISPR-Cas on conjugative antibiotic resistance dissemination in a mammalian intestinal model. We discovered that CRISPR-Cas is a strikingly robust barrier to pAM714 and pAM771 acquisition in the murine intestine.

## Results

### Mating conditions impact CRISPR-Cas activity against pAM714

We analyzed planktonic and agar plate mating reactions between *E. faecalis* OG1SSp(pAM714) donors and T11RF or T11RF Δ*cas9* recipients over an 18-hour period (Fig. 1; see Table 1 for strain details). We inoculated mating reactions at donor to recipient ratios of 1:9 and 1:1 (Fig. 1). Donors were quantified by plating matings on media with spectinomycin, streptomycin, and erythromycin (Fig. S1); transconjugants with rifampicin, fusidic acid, and erythromycin (Fig. 1), and total recipients (which includes transconjugants) with rifampicin and fusidic acid (Fig. 2-3). In our experiments, we used erythromycin resistance to track pAM714 conjugation. *ermB* is encoded on Tn*917*, which theoretically could transpose from pAM714 into the *E. faecalis* chromosome, thereby unlinking erythromycin resistance from pAM714 presence. However, Tn*917* transposition frequencies are very low (10^-6^) in the absence of the inducer erythromycin (31). No mating reactions in our study contained erythromycin.

**Figure 1.**
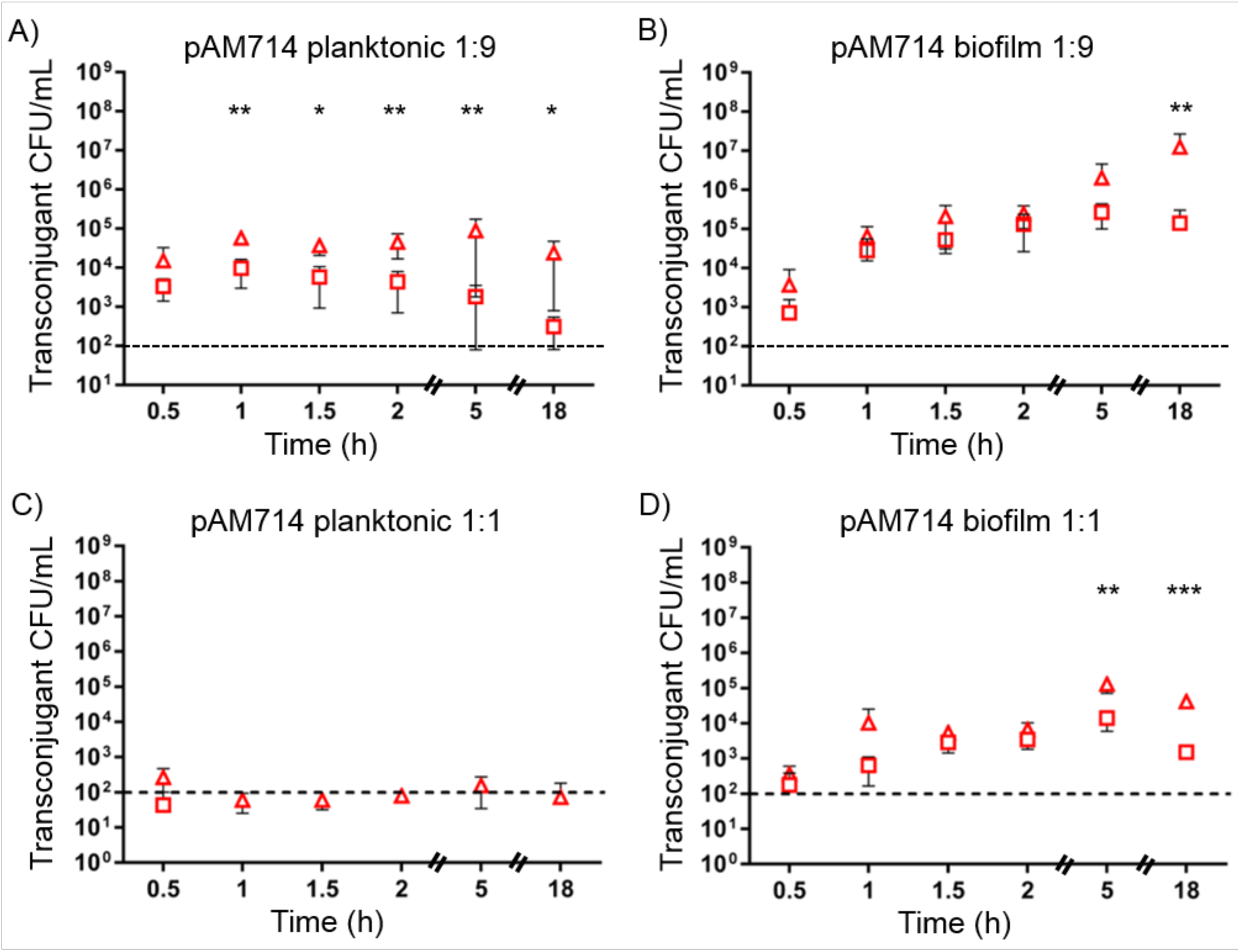
Impact of CRISPR-Cas on pAM714 transconjugant yields under different *in vitro* conditions. The CFU/mL of transconjugants obtained in mating reactions sampled over an 18-hour period is shown for T11RF (squares) and T11RFΔ*cas9* (triangles) recipient strains. Conjugation was performed under planktonic conditions in broth (A and C) and biofilm conditions on an agar plate (B and D) utilizing OG1SSp as a donor strain. Conjugation reactions were initiated with a 1:9 (A and B) or 1:1 (C and D) donor to recipient ratio. The limit of detection is indicated by the dashed line. Data shown are the average and standard deviation from a minimum of three independent trials for each time point. Statistical significance was assessed using a two-tailed Student’s t-Test; *P*-values, *<0.05, **<0.01 and ***<0.001.

**Figure 2.**
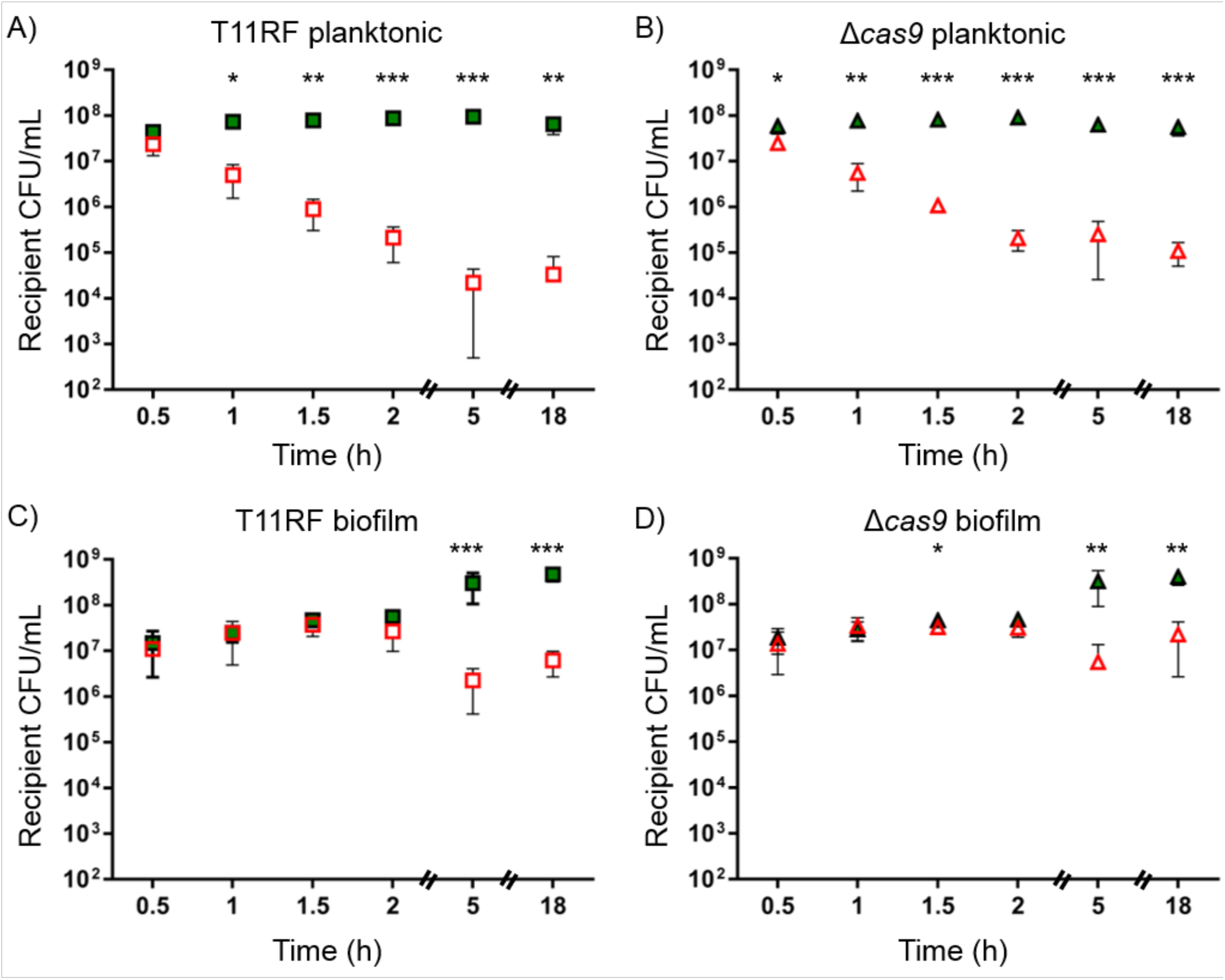
Recipient cell densities for *in vitro* conjugations at a 1:9 donor to recipient ratio. T11RF (squares) and T11RFΔ*cas9* (triangles) recipient cell densities in CFU/mL was determined for both planktonic (A and B) and biofilm (C and D) mating reactions with pAM714 (open, red symbols) and pAM771 (closed green symbols) donors. The limit of detection was 10^2^ CFU/mL. Data shown are the average and standard deviation from a minimum of three independent trials for each time point for both mating conditions. Statistical significance was assessed using a two-tailed Student’s t-Test; *P*-values, *<0.05, **<0.01 and ***<0.001.

**Figure 3.**
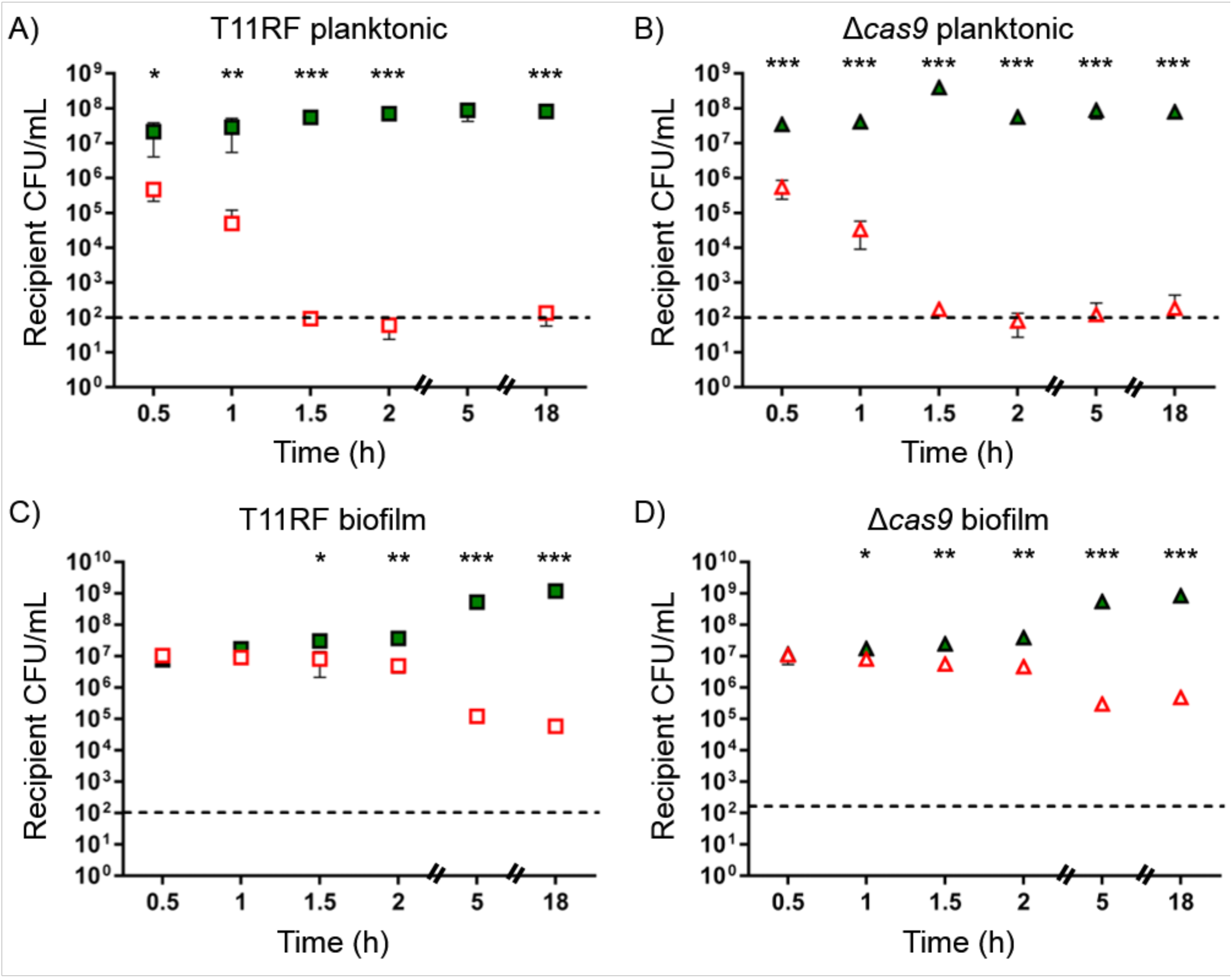
Recipient cell densities for *in vitro* conjugations at a 1:1 donor to recipient ratio. T11RF (squares) and T11RFΔ*cas9* (triangles) recipient cell densities in CFU/mL was determined for both planktonic (A and B) and biofilm (C and D) mating reactions with pAM714 (open, red symbols) and pAM771 (closed green symbols) donors. The limit of detection is indicated by the dashed line. Data shown are the average and standard deviation from a minimum of three independent trials for each time point for both mating conditions. Statistical significance was assessed using a two-tailed Student’s t-Test; *P*-values, *<0.05, **<0.01 and ***<0.001.

**Table 1.** *E. faecalis* strains used in this study.

| Strain Name | Description | Reference |
| --- | --- | --- |
| T11RF | Rifampicin-fusidic acid resistant derivative of strain T11 | (19, 37) |
| T11RFΔ*cas9* | T11RF with an in-frame deletion of *cas9* | (19) |
| OG1SSp(pAM714) | Spectinomycin-streptomycin resistant derivative of strain OG1 harboring pAM714, conferring erythromycin resistance via Tn917 insertion upstream of the *par* locus; *cyl*<sup>+</sup> | (25, 26) |
| OG1SSp(pAM771) | Spectinomycin-streptomycin resistant derivative of strain OG1 harboring pAM771, conferring erythromycin resistance via Tn917 insertion disrupting *cylL* of the cytolysin operon; *cyl*<sup>-</sup> | (25, 29, 30) |

For 1:9 donor:recipient ratio experiments, we observed ∼10^3^-10^4^ transconjugants for both T11RF and Δ*cas9* recipients after 30 minutes of mating (Fig. 1A-B). T11Δ*cas9*(pAM714) transconjugant numbers remained stable for the remainder of the planktonic mating experiment, while T11RF(pAM714) transconjugant numbers decreased (Fig. 1A). In contrast, pAM714 transconjugant yields in biofilm matings rose over time for both T11RF and Δ*cas9* recipients, up to the 2 hour time point. After that, T11RF(pAM714) transconjugant numbers did not increase further, while T11Δ*cas9*(pAM714) transconjugants increased by 2 log (Fig. 1B). For both planktonic and biofilm matings, we observed significant differences in transconjugant yields between T11RF and Δ*cas9* recipients at the experiment end point (18 hours) and for some earlier time points. We note that, despite CRISPR-Cas activity, ∼10^5^ pAM714 transconjugants were still observed for T11RF recipients in biofilms (Fig. 1B).

We next assessed conjugation using an equal (1:1) donor:recipient ratio. Increasing donor densities relative to recipients reduces pheromone detection by pheromone-responsive plasmids (32). Transconjugant numbers were overall lower than those observed for 1:9 ratio experiments (Fig. 1C-D). For planktonic matings, transconjugant numbers were at or below our limit of detection, therefore the impact of *cas9* on transconjugant yield could not be assessed (Fig. 1C). Transconjugants were detected for biofilm matings (Fig. 1D), but the yields were lower than those observed for 1:9 ratio experiments (Fig. 1B). Nevertheless, *cas9* protected recipients from pAM714 acquisition at the 5 and 18 h time points (Fig. 1D).

### Cytolysin activity depletes recipient cells irrespective of functional CRISPR-Cas

We hypothesized that the cytolysin encoded by pAM714 could kill recipient cells that utilize CRISPR-Cas against the plasmid. pAM771 is isogenic with pAM714, except that the Tn*917* insertion disrupts *cylL* of the cytolysin biosynthesis gene cluster (25, 29, 30). pAM714 and pAM771 have been utilized in previous studies assessing the impact of cytolysin on virulence, hamster intestinal colonization, and plasmid transfer (29, 30, 33). We performed planktonic and biofilm mating reactions with OG1SSp(pAM771) donors and compared the results with the OG1SSp(pAM714) mating experiments.

Recipient (Fig. 2-3) but not donor (Fig. S1-S2) densities were substantially impacted in all pAM714 mating reactions, irrespective of *cas9* presence or absence. The effect was stronger in planktonic matings (Fig. 2A-B and Fig. 3A-B) than in biofilm matings (Fig. 2C-D and Fig. 3C-D), and strongest in planktonic matings at a 1:1 donor:recipient ratio, where recipient numbers fell to below the limit of detection after 1.5 h of mating (Fig. 3A-B). These results are consistent with pAM714 transconjugant yields under these conditions (Fig. 1C). In biofilm matings, striking effects on recipient cell densities were not observed until later time points (5 h and 18 h; Fig. 2C-D and Fig. 3C-D).

Unlike observations from pAM714 matings, recipient numbers were stably high in pAM771 matings. Moreover, pAM771 transconjugant yields were not substantially impacted by donor:recipient ratio (Fig. 4). Similar transconjugant yields were detected for planktonic matings at 1:9 (Fig. 4A) and 1:1 (Fig. 4C) ratios, and for biofilm matings at the two ratios (Fig. 4B and Fig. 4D, respectively). The effect of *cas9* was minor in magnitude but statistically significant at the end of planktonic mating. Deletion of *cas9* increased plasmid acquisition significantly, by ∼2 log, after 18 h of biofilm mating.

**Figure 4.**
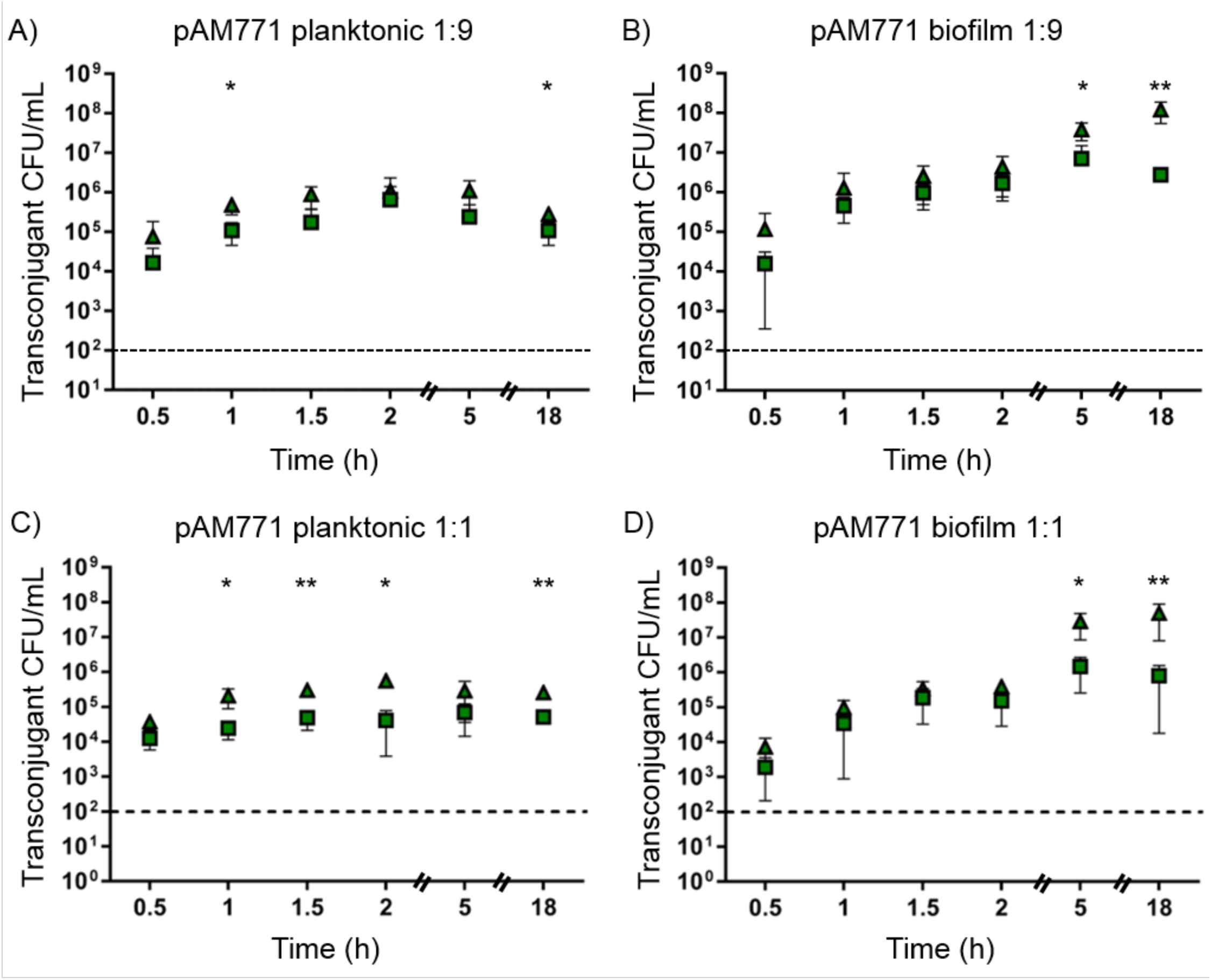
Impact of CRISPR-Cas on pAM771 transconjugant yields under different *in vitro* conditions. The CFU/mL of transconjugants obtained in mating reactions sampled over an 18-hour period is shown for T11RF (squares) and T11RFΔ*cas9* (triangles) recipient strains. Conjugation was performed under planktonic conditions in broth (A and C) and biofilm conditions on an agar plate (B and D) utilizing OG1SSp as a donor strain. Conjugation reactions were initiated with a 1:9 (A and B) or 1:1 (C and D) donor to recipient ratio. The limit of detection is indicated by the dashed line. Data shown are the average and standard deviation from a minimum of three independent trials for each time point. Statistical significance was assessed using a two-tailed Student’s t-Test; *P*-values, *<0.05, **<0.01 and ***<0.001.

Our results overall with *in vitro* experiments demonstrate that planktonic versus biofilm settings, different donor/recipient ratios, production of a plasmid-encoded bacteriocin, and the time points at which matings are sampled all impact transconjugant yields and conclusions reached about the apparent activity of CRISPR-Cas. Moreover, CRISPR tolerance is consistently observed *in vitro*, with the exception of settings where little plasmid transfer occurs into any recipient (pAM714 planktonic matings at a 1:1 donor:recipient ratio; Fig. 1C).

### *cas9* expression in T11RF biofilms and planktonic cultures

Little is known about the transcriptional or post-transcriptional regulation of *cas9* in *E. faecalis*. During our investigation of a different Type II CRISPR-Cas system in *E. faecalis*, CRISPR1-Cas, we observed that 27-fold overexpression of *cas9* enhanced *in vitro* defense against PRP-mobilizable plasmids by ∼2-3 logs (27). Because low *cas9* expression in wild-type T11RF under *in vitro* conditions could impact CRISPR-Cas defense, we attempted to assess T11RF *cas9* expression levels under these conditions.

We first assessed T11RF *cas9* expression in monoculture under the same planktonic and biofilm conditions used for mating (Fig. 5). We normalized to *recA* because this was the normalization used for our previous assessment of engineered *cas9* expression in a different *E. faecalis* strain (27). We observed very low expression of *cas9* relative to *recA* in the early biofilm time points, and a dramatic increase in this ratio at the 5 and 18 hour time points (Fig. 5B). While this is consistent with the impact of *cas9* on transconjugant yields in biofilm mating experiments (Fig. 1 and Fig. 4), an important caveat is that we noted substantially lower *recA* expression at 5 h and 18 h in biofilm T11RF cultures relative to earlier time points (Dataset S1), and thus *cas9* levels may be under represented at these later time points. For planktonic cultures, relative *cas9* expression levels varied in the mid-range between the high and low extremes observed during biofilm growth (Fig. 5A). This, too, is consistent with transconjugant yields observed in planktonic settings, where a robust effect of *cas9* was not observed for pAM771 matings (Fig. 4), and in pAM714 matings the killing of recipient cells was a major confounder (Fig. 2-3). We did not observe the same variation over time in *recA* expression in planktonic cultures as we did for biofilm cultures (Dataset S1). Overall, the relative expression of *cas9* to *recA* is consistent with *cas9* impact on transconjugant yields, but the mechanism for this is unclear.

**Figure 5.**
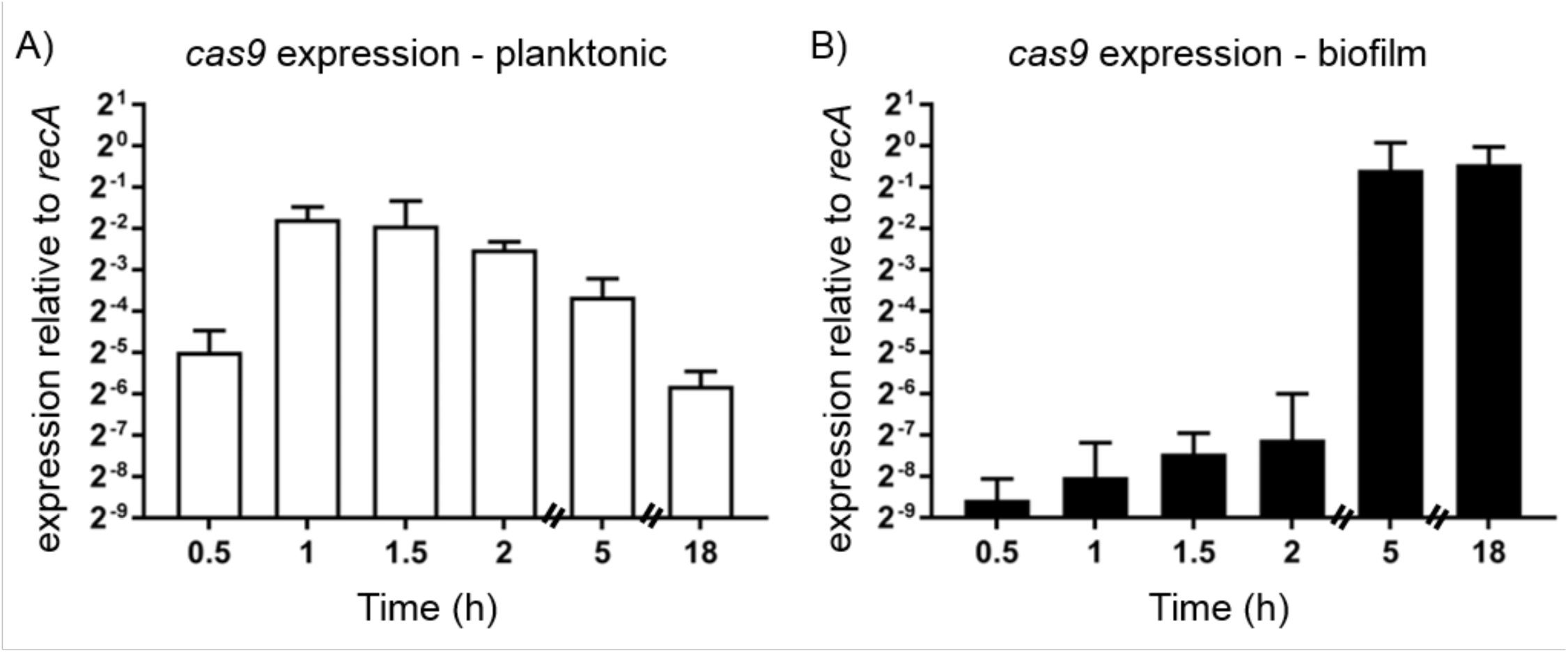
T11RF *cas9* expression in planktonic culture and agar plate biofilms. Expression of *cas9* was assessed by RT-qPCR under identical growth conditions used for planktonic (A) and biofilm (B) conjugation experiments, with the exception that the OG1SSp plasmid donor strain was not present. Expression of *cas9* is expressed relative to *recA*. Data shown represent the average *cas9* expression from three independent experiments.

### CRISPR-Cas is a robust barrier to PRP acquisition in the murine intestine

We assessed CRISPR3-Cas activity against pAM714 and pAM771 in a mouse model of *E. faecalis* intestinal dysbiosis. To establish antibiotic-induced dysbiosis, mice were administered a cocktail of antibiotics in their drinking water for seven days, followed by placement on regular water for 24 h. The mice were colonized sequentially with recipient and donor *E. faecalis* strains at a 1:1 donor:recipient ratio. Fecal pellets were collected at 24, 48, and 96 hours post co-colonization, homogenized, and the number of transconjugants, donors, and recipients were quantified (Fig. 6). Experimental groups consisting of different combinations of donor and recipient strains were used: OG1SSp with T11RF as a plasmid-free control group, OG1SSp(pAM714) donors with T11RF recipients, and OG1SSp(pAM714) donors with T11RFΔ*cas9* recipients. In separate experiments, OG1SSp(pAM771) donors were used.

**Figure 6.**
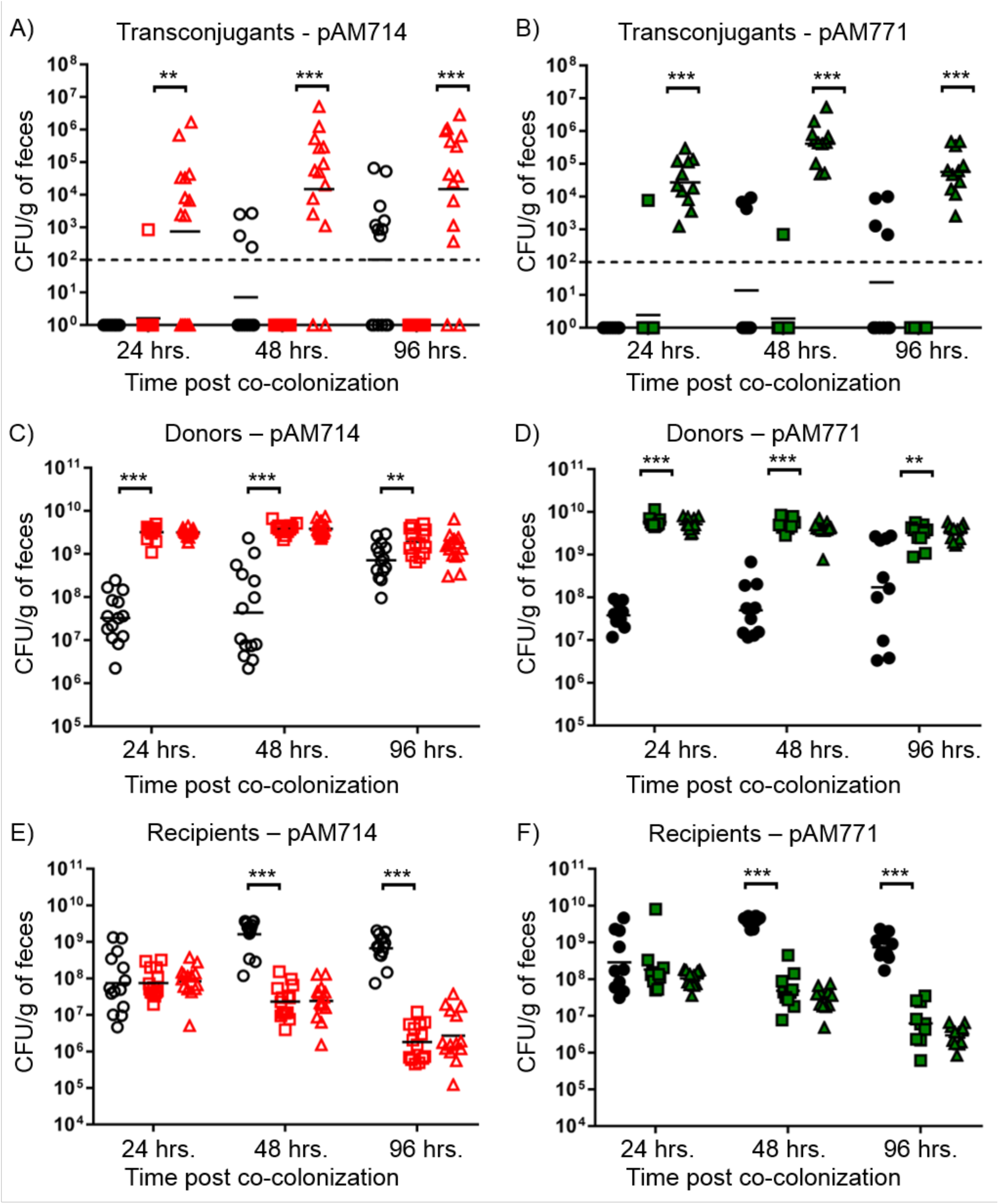
Impact of CRISPR-Cas on plasmid transfer in the mouse intestine. The number of transconjugant (A, B), donor (C, D), and recipient (E, F) CFU/g of feces for individual mice were determined by plating feces on selective agars. Each symbol represents one mouse. Experimental groups are described in the materials and methods. Black horizontal bars represent the geometric mean of data in each group. Shown are data for *in vivo* pAM714 (A, C, and E; open red symbols) pAM771 (B, D, and F; closed green symbols) transfer. Control mice co-colonized by OG1SSp and T11RF are represented by open circles. Mice colonized with T11RF recipients are represented by squares. Mice colonized with T11RFΔ*cas9* recipients are represented by triangles. The limit of detection is indicated by the dashed line. Statistical significance was assessed using a two-tailed Student’s t-Test; *P*-value, **<0.01 and ***<0.001.

We detected pAM714 transconjugants in only one of fourteen mice colonized with T11RF recipients at 24 h post co-colonization, and for none of the mice at 48 and 96 h time points (Fig 6A). Strikingly, pAM714 transconjugants at densities up to ∼5 x 10^6^ CFU/g of feces were observed for twelve of fourteen mice colonized with T11RFΔ*cas9* recipients over the course of the experiment (Fig. 6A). We screened 36 presumptive T11RFΔ*cas9*(pAM714) transconjugants by PCR amplification of the pAM714 *repB* gene; all possessed this gene (Fig. S3). We observed that some control (no plasmid) mice at 48 and 96 h time points had colony growth on media with selection for transconjugants (i.e. media supplemented with rifampicin, fusidic acid, and erythromycin) (Fig. 6A). We screened 20 of these colonies by PCR amplification of the pAM714 *repB* gene; none possessed this gene (Fig. S3). We infer that recipients received erythromycin resistance determinants from the native mouse microbiota via a non-pAM714-dependent mechanism.

We performed identical *in vivo* conjugation experiments with OG1SSp(pAM771) donors. Fewer mice were observed with sporadic erythromycin resistance in the control group for pAM771 experiments (Fig. 6B). T11RF(pAM771) transconjugants were detected for only one of ten mice at each of the 24 and 48 h time points, whereas T11RFΔ*cas9*(pAM771) transconjugants were detected in all eleven mice and at all time points (Fig. 6B).

Overall, these data demonstrate that there is a profound impact of CRISPR-Cas on plasmid transfer between *E. faecalis* strains in the dysbiotic mouse intestine. These observations are in contrast to any *in vitro* condition evaluated, where either plasmid transfer was not observed (for 1:1 ratio planktonic matings) or transconjugants arose despite recipients having CRISPR-Cas defense. Moreover, cytolysin did not impact *in vivo* plasmid transfer, as was observed for *in vitro* transfer. This is consistent with a previous study that analyzed transfer of pAM714 and pAM771 between *E. faecalis* in the hamster intestinal tract (29).

### Potential cytolysin-independent *in vivo* colonization benefit to strains possessing a PRP

We next assessed whether cytolysin impacted the colonization of *E. faecalis* donors in the mouse intestine. We compared donor densities in control mice colonized with OG1SSp to those colonized with OG1SSp(pAM714) (Fig. 6C) or OG1SSp(pAM771) (Fig. 6D). We observed no benefit to donors possessing pAM714 versus pAM771. However, donor densities in the control group were significantly reduced compared to plasmid-bearing donors at all time points. These data suggest that there is a cytolysin-independent colonization benefit for OG1SSp harboring pAM714 or pAM771. This is consistent with recent observations for *E. faecalis* harboring the PRP pCF10 during intestinal colonization of germ-free mice (34).

At 24 hours post-co-colonization, recipient strain densities from control, pAM714, and pAM771 test groups were similar (Fig. 6E-F). On average, T11RF recipient densities increased in control mice at subsequent time points, but decreased in both pAM714 and pAM771 test groups (Fig. 6E-F). No differences were observed for recipient densities in pAM714 versus pAM771 groups. This demonstrates that the reduction in recipient cell densities observed *in vitro* for pAM714 but not pAM771 matings (Fig. 2-3) does not occur in the *in vivo* model tested here. Rather, our data suggest that there is a cytolysin-independent fitness advantage for pAM714/pAM771 donors *in vivo*. We also note that no differences were observed in T11RF and T11RFΔ*cas9* colonization, indicating that the presence or absence of *cas9* does not impact intestinal colonization success in this model (Fig. 6E-F).

## Discussion

We have found that native CRISPR-Cas encoded by a member of the mammalian intestinal microbiota can block the *in vivo* dissemination of an antibiotic resistance plasmid in a murine intestinal colonization model. This is in contrast to *in vitro* observations, where the same plasmid is frequently acquired by recipient cells despite CRISPR-Cas. For the *E. faecalis* CRISPR1-Cas system we previously investigated (27, 28), these tolerant cells harboring both CRISPR-Cas and a plasmid it targets have an *in vitro* growth defect that is resolved by either plasmid loss or by mutation of CRISPR-Cas when antibiotic selection for the plasmid is applied. We did not detect CRISPR tolerance *in vivo*. One possible explanation for this is that CRISPR-Cas is far more effective *in vivo* than *in vitro*, and transconjugants never arise in cells possessing functional CRISPR-Cas *in vivo*. Another is that they do arise, but their growth defect combined with turnover of intestinal contents results in their rapid elimination *in vivo*. One method to test this would be to add erythromycin selection *in vivo*; we would expect to observe high densities of T11RF plasmid transconjugants that are CRISPR-Cas mutants. We were not able to test this in our current model system because of the erythromycin inducibility of Tn*917*, which would complicate plasmid detection.

What mechanisms underlie our observations about the impact of CRISPR-Cas on conjugative plasmid transfer *in vitro* and *in vivo*? Several factors may factor in this process, including plasmid host range, donor to recipient ratios and their relative colonization densities, community spatial structure (i.e. biofilms), flow and dilution rate, nutrient availability, community diversity and the relative densities of plasmid-susceptible versus non-susceptible hosts, and selection for the plasmid. With the PRPs, there is the additional consideration of pheromone concentration; the pheromone is a short peptide elaborated by recipient *E. faecalis* cells (and some other bacteria) that induces transcription of conjugation genes in the donor strain (11, 12). Finally, there are CRISPR-Cas-specific factors about which little is known, such as the *in vitro* versus *in vivo* transcriptional and post-transcriptional regulation of *cas9*.

We confirmed that several of these factors influence PRP transconjugant yield *in vitro*. *In vitro cas9* expression appears to be insufficient to confer protection to many *E. faecalis* recipients. Other factors having strong effects were cytolysin biosynthesis encoded by the plasmid, which negatively impacted recipient densities, and the donor:recipient ratio, which affects induction of conjugation by pheromone signaling (32).

We determined that the pAM714/pAM771 conjugation frequency to T11RFΔ*cas9* recipients is ∼10^0^-10^-2^ transconjugants per donor (TC/D) for *in vitro* broth and agar plate biofilm experiments, while *in vivo* it ranges from 10^-3^ to 10^-7^ (Fig. S4). Conjugation frequency of PRPs can be modulated by deleting aggregation substance in the plasmid, which should reduce conjugation frequency in broth cultures but not biofilms (11), or by interfering with pheromone production by plasmid-free recipient cells. This will be addressed in future work. Also to be addressed is the effect of varying the total cell count of the donors and recipients at the time of culture inoculation; initiating cultures with fewer cells may more accurately reflect the nature of intestinal colonization by *E. faecalis*.

In the *in vivo* model used here, we induced intestinal dysbiosis with antibiotics, allowed mice to recover for one day, and then colonized them with *E. faecalis*. This models what can occur in patients after receiving antibiotic therapy. Another mouse model used in the field establishes long-term colonization of *E. faecalis* without major disruption of normal intestinal microbiota (35). Further, a recent study utilized a germ-free mouse model to examine *in vivo* transfer of the PRP pCF10 among intestinal *E. faecalis* (34). In the germ-free model, enterococci achieve very high densities and diversity is very low. In the native colonization model, diversity is high, and production of the Bac-21 bacteriocin from the PRP pPD1 significantly enhances *E. faecalis* colonization (35). These two models can be used to assess how community diversity and the densities of plasmid-susceptible and non-susceptible hosts impact CRISPR-Cas efficacy *in vivo*.

How far can we extrapolate from studies with *E. faecalis* to other members of the mammalian microbiota, and from PRPs to other plasmids with different properties and host ranges? Put another way, does CRISPR-Cas encoded by other members of the native microbiota confer the same robust defense against antibiotic resistance plasmids as observed for *E. faecalis* and PRPs? Will *E. faecalis* CRISPR-Cas defense against non-PRP plasmids be equally robust? Much future work remains to elucidate these questions.

## Materials and Methods

### Bacteria and reagents used

Strains used in this study are shown in Table 1. *E. faecalis* strains were cultured in brain heart infusion (BHI) broth or on BHI agar at 37°C. Antibiotic concentrations used were as follows: rifampicin, 50 µg/mL; fusidic acid, 25 µg/mL; spectinomycin, 500 µg/mL; streptomycin, 500 µg/mL; erythromycin, 50 µg/mL. Antibiotics were purchased from Sigma-Aldrich or Research Products International (RPI).

### Conjugation experiments

Donor and recipient strains were cultured overnight in BHI broth in the absence of antibiotic selection. The following day, cultures were diluted 1:10 into fresh BHI and incubated at 37°C for 1.5 hours. For planktonic conjugations at a 1:9 donor:recipient ratio, 2 mL of donor and 18 mL of recipient were mixed in a flask and incubated without agitation at 37°C for 30 min to 18 h. For planktonic conjugations at a 1:1 donor:recipient ratio, 10 mL of donor and 10 mL of recipient were mixed in a flask and incubated without agitation at 37°C for 30 min to 18 h. At each time point, 1 mL of the mating reaction was removed and used for serial dilutions and plating on selective media. For biofilm mating reactions at a 1:9 donor:recipient ratio, 100 µL of donor was mixed with 900 µL of recipient, and for reactions at a 1:1 donor:recipient ratio, 500 µL of donor was mixed with 500 µL of recipient. The mixture was centrifuged for 1 min at 16,000 x g. After centrifugation, 100 µL supernatant was used to resuspend the pellet, which was then spread-plated on non-selective BHI agar. To allow for sampling of multiple time points of biofilms, multiple identical conjugation reactions were generated using the same donor and recipient inocula. The conjugation reactions were incubated at 37°C for 30 min to 18 h. At each time point, cells were collected by washing and scraping an agar plate using 2 mL 1X phosphate buffered saline (PBS) supplemented with 2 mM EDTA, and serial dilutions were plated on selective media. For all matings, BHI agar supplemented with antibiotics was used to quantify the donor (spectinomycin, streptomycin, and erythromycin), recipient (rifampicin and fusidic acid), and transconjugant (rifampicin, fusidic acid, and erythromycin) populations. Plates were incubated for 36-48 h at 37°C. Plates with 30 to 300 colonies were used to calculate CFU/mL.

### Mouse model of *E. faecalis* colonization

Seven days prior to bacterial colonization, 6-8 week old C57BL/6 mice were gavaged with 100 µL of an antibiotic cocktail (streptomycin 1 mg/mL, gentamicin 1 mg/mL, erythromycin 200 µg/mL), and given a water bottle ad libitum with the same antibiotic cocktail for 6 days following gavage. 24 h prior to bacterial inoculation, antibiotic water was removed and replaced with standard sterile antibiotic-free water. Bacteria were grown overnight in BHI, and mice were gavaged with 10^9^ CFU in PBS of each bacterial strain as experimental groups indicated (1:1 donor:recipient ratio). Samples used for gavage were plated on BHI to confirm that inocula were equal across strains. Fecal samples from mice were collected at 0 h, 24 h, 48 h and 96 h. Fecal samples were resuspended in 1 mL of sterile PBS and dilutions were plated on BHI agar supplemented with antibiotics to quantify the donor (spectinomycin, streptomycin, and erythromycin), recipient (rifampicin and fusidic acid), and transconjugant (rifampicin, fusidic acid, and erythromycin) populations. Plates were incubated for 36-48 h at 37°C. Plates with 30 to 300 colonies were used to calculate CFU/g of feces. Experiments were performed in duplicate or triplicate as follows: For OG1SSp pAM714/T11RF(+/-*cas9*) co-colonization, three independent experiments were performed consisting of 4, 4, and 6 mice per group per experiment. For OG1SSp pAM771/T11RF(+/-*cas9*) co-colonization, two independent experiments were performed consisting of 5 mice per group, except in the second experiment where 5 mice were used for the control and wild-type T11RF groups and 6 mice were used for the T11RF Δ*cas9* group. Data from individual experimental replicates were combined and graphed together. All animal protocols were approved by the Institutional Animal Care and Use Committee of the University of Colorado Anschutz Medical Campus (protocol number 00253).

### Colony PCR to verify *in vivo* transconjugants

Fecal pellets were collected at 0 hr, 24 hr, 48 hr and 96 hr, weighed, and resuspended in 1 mL PBS. 20 µL were plated at multiple dilutions on BHI containing rifampicin, fusidic acid, and erythromycin. Individual colonies were picked, resuspended in 20 µL nuclease-free water, and 1 µL used in PCR with Taq DNA Polymerase (New England Biolabs). Primers amplified the *repB* region of plasmids pAM714 and pAM771 (pAD1 *repB*-For: 5’-CGT TCC ATG TGT CTA ACA ATT GTA TTA AAC-3’ and pAD1 *repB*-Rev: 5’-CGA TGA TGG TAG CAA TTC AAG AAG G-3’).

### *In vitro* T11RF *cas9* expression analysis

Identical growth conditions and procedures were used as described above for planktonic and biofilm conjugation experiments with the exception that the OG1SSp plasmid donor strain was not present. At the desired time point, a 5 mL culture aliquot was collected from planktonic cultures conditions, or biofilms were collected from agar plates using 2 mL of PBS supplemented with 2 mM EDTA. Total RNA was isolated using RNA-Bee and chloroform precipitation as previously described (36). 200 ng RNA was used as template for cDNA synthesis using qScript cDNA Supermix (Quanta Biosiences). Subsequent qPCR reactions were performed using the AzuraQuant Green Fast qPCR Mix LoRox (Azura Genomics). Primer sequences to query *cas9* were *cas9*-For: 5’-GCA ACT GGG ATG ACT ATC A-3’ and *cas9*-Rev: 5’-GCA TAA CGC GTA TCA TTC A-3’. Primer sequences to query *recA* were *recA*-For: 5’-TGG TGA GAT GGG AGC GAG CC-3’ and *recA*-Rev: 5’-TCA GGA TTT CCG AAC ATC ACG CC-3’. Expression of *cas9* relative to *recA* was calculated as 2ΔCq, where ΔCq = (CqT11RF @ time x *cas9* – CqT11RF @ time x *recA*). Data shown represents the average *cas9* expression from three independent experiments for both growth conditions assessed.

## Acknowledgments

The authors thank Ian Jorgeson with assistance with figure formatting and Dr. Michael Gilmore for providing pAM714 and pAM771. This work was supported by grants R01AI116610 to KLP, R01AI141479 to BAD, and K01DK102436 to BAD from the National Institutes of Health. The authors declare that the funders of this work had no role in the design of experiments, interpretation of data, or the decision to publish this work.

## Supplemental figure legends

**Figure S1.**
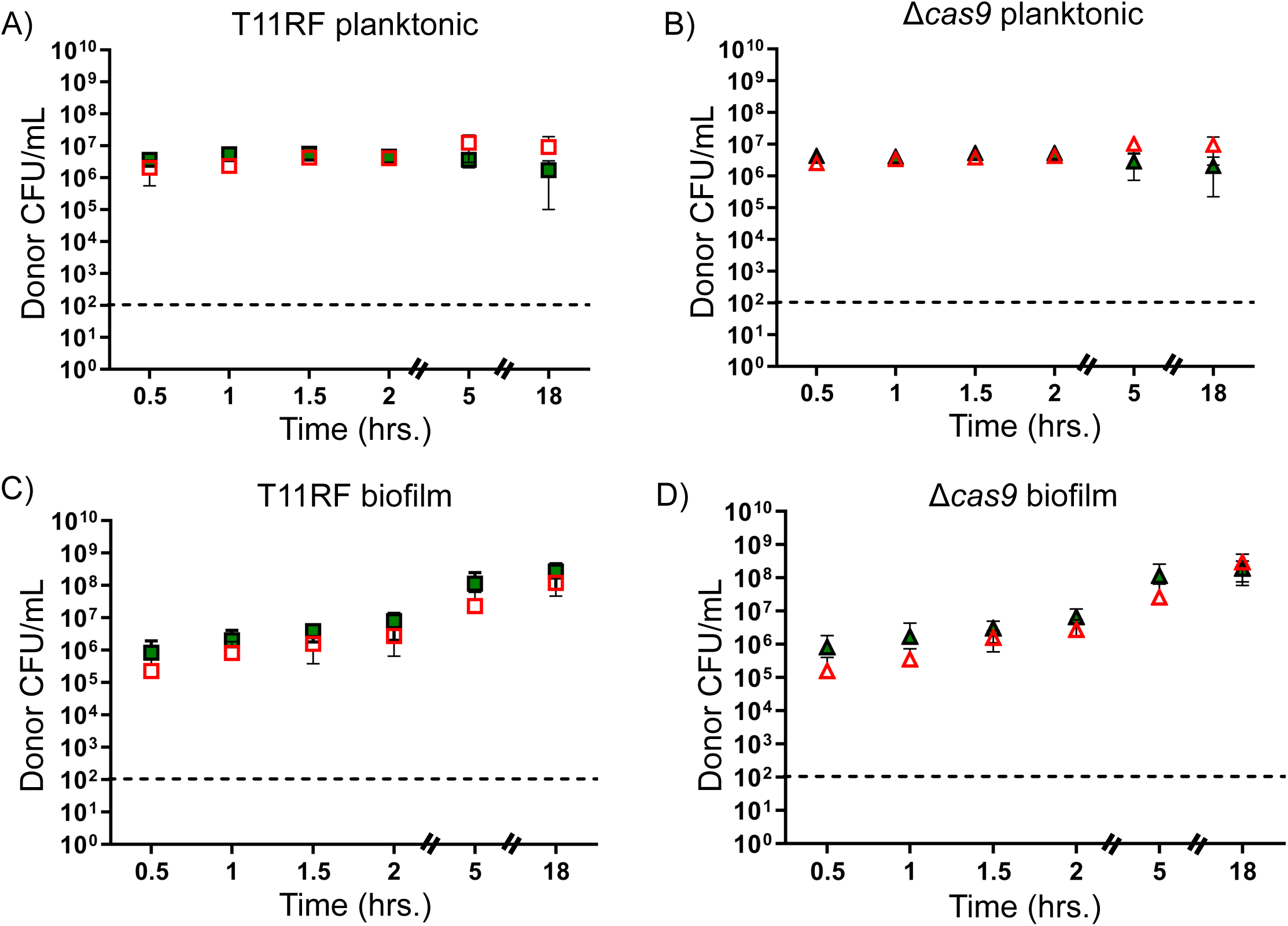
Donor cell densities for *in vitro* conjugations at a 1:9 donor to recipient ratio. T11RF (squares) and T11RFΔ*cas9* (triangles) donor cell densities in CFU/mL was determined for both planktonic (A and B) and biofilm (C and D) mating reactions with pAM714 (open, red symbols) and pAM771 (closed green symbols) donors. The limit of detection was 10^2^ CFU/mL. Data shown are the average and standard deviation from a minimum of three independent trials for each time point for both mating conditions. Statistical significance was assessed using a two-tailed Student’s t-Test; *P*-values, *<0.05, **<0.01 and ***<0.001.

**Figure S2.**
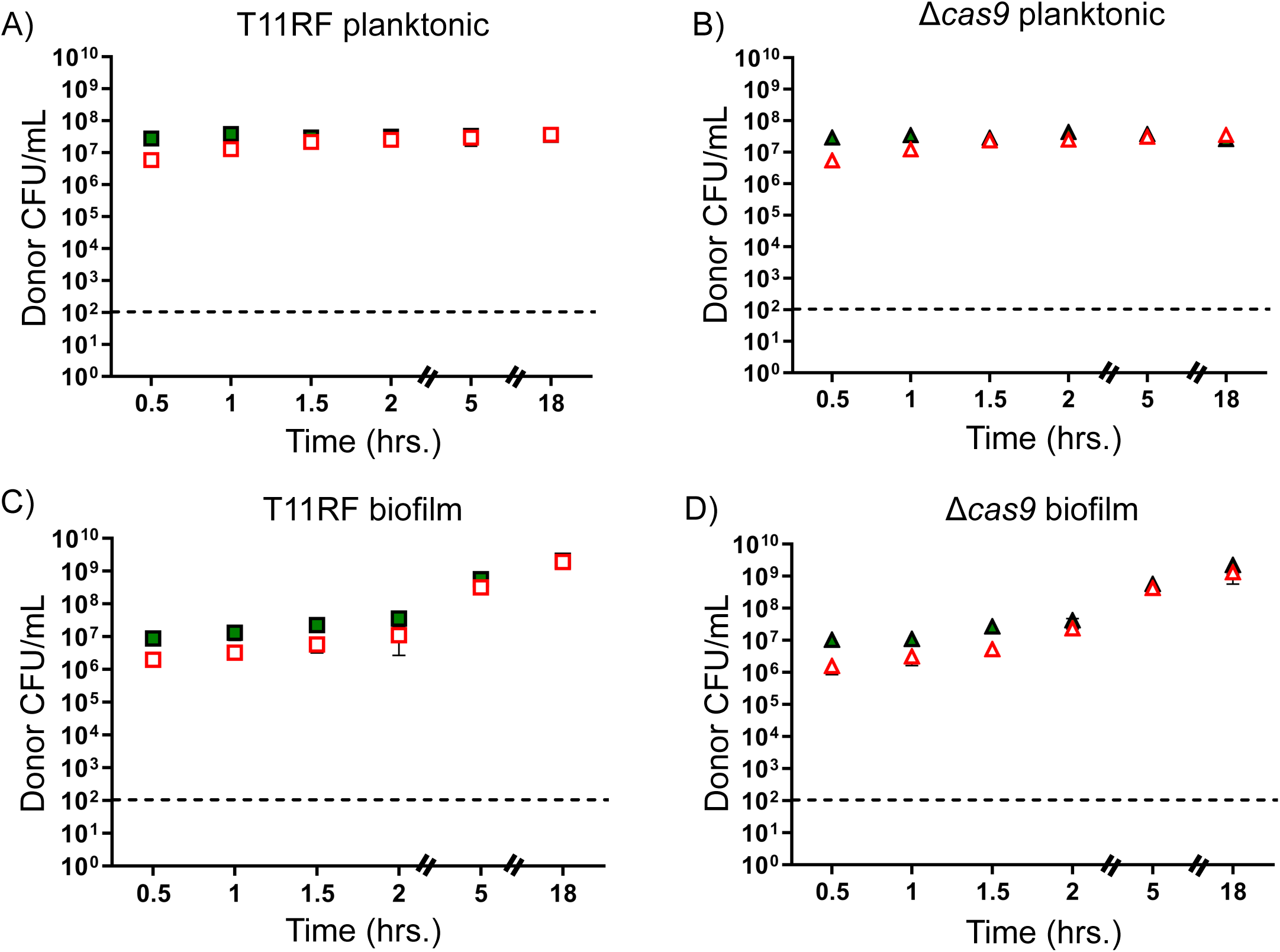
Donor cell densities for *in vitro* conjugations at a 1:1 donor to recipient ratio. T11RF (squares) and T11RFΔ*cas9* (triangles) donor cell densities in CFU/mL was determined for both planktonic (A and B) and biofilm (C and D) mating reactions with pAM714 (open, red symbols) and pAM771 (closed green symbols) donors. The limit of detection was 10^2^ CFU/mL. Data shown are the average and standard deviation from a minimum of three independent trials for each time point for both mating conditions. Statistical significance was assessed using a two-tailed Student’s t-Test; *P*-values, *<0.05, **<0.01 and ***<0.001.

**Figure S3.**
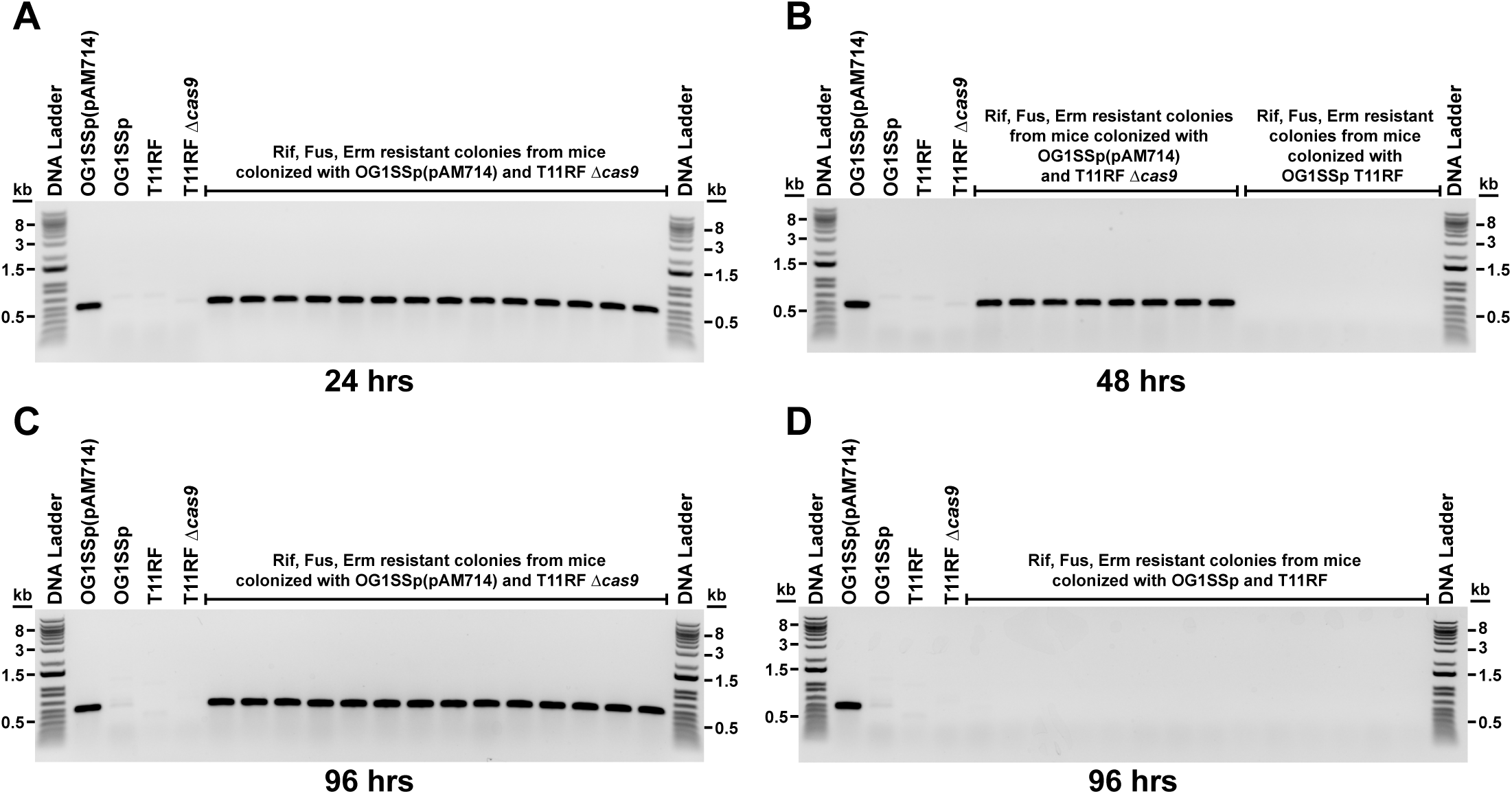
PCR confirms that experimental transconjugants carry pAM714 and spontaneous resistant isolates from control animals do not. Agarose gels show PCR amplification products for the *repB* gene of pAM714 at 24 (A), 48 (B) and 96 (C) hours post-colonization, and the lack of a *repB* amplification signal in isolates originating from control mice and growing on transconjugant selection agar (B and D). PCR reactions for the presence and absence of the *repB* gene using strains OG1SSp(pAM714), OG1SSp, T11RF and T11RF Δ*cas9* are included on each gel. Rif – Rifampicin, Fus – Fusidic acid, Erm – Erythromycin.

**Figure S4.**
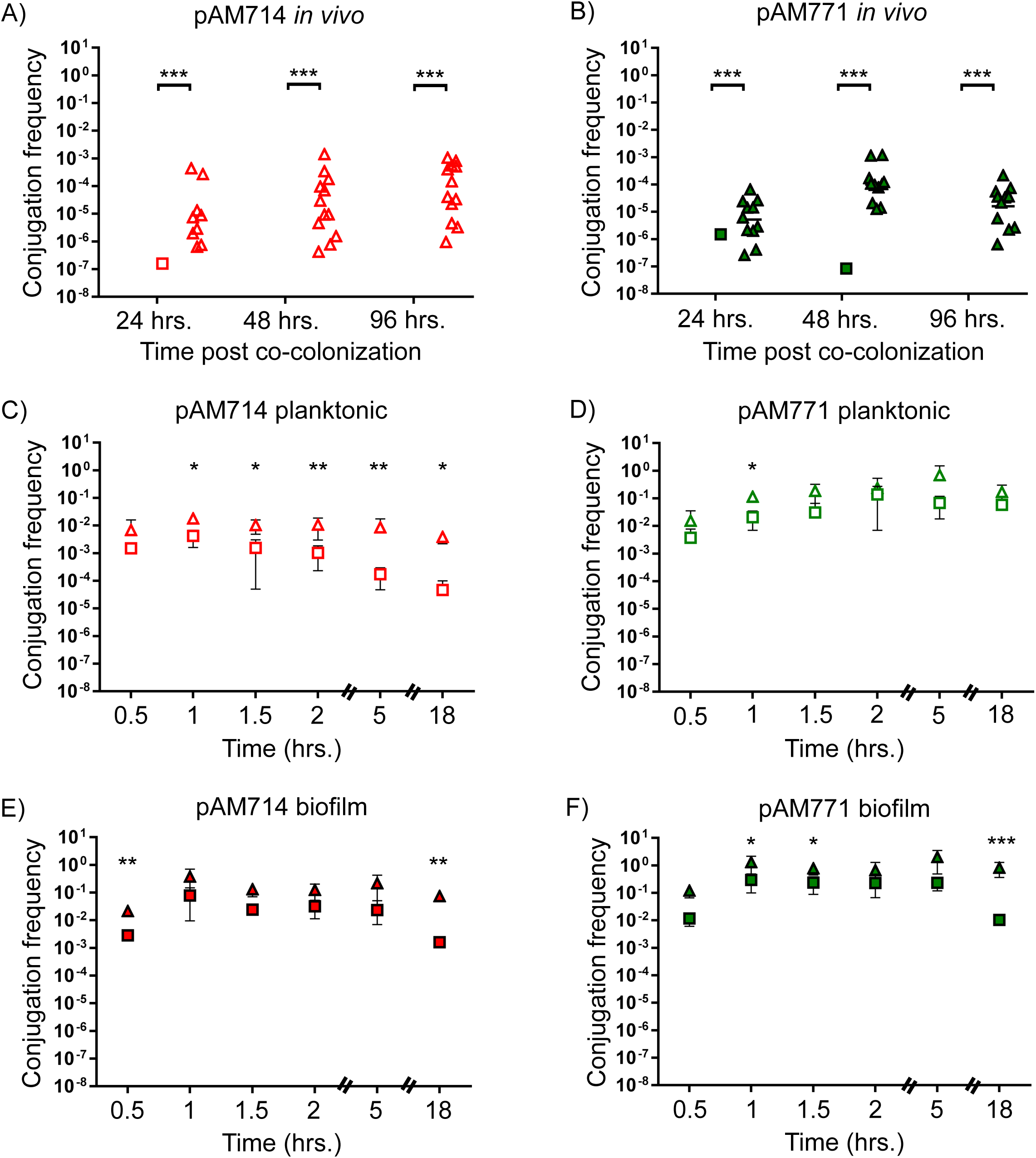
Frequency of conjugation *in vitro* and in the mouse intestine. Conjugation frequencies for pAM714 under mouse intestine (A), planktonic (C), and biofilm (E) settings are shown as transconjugants per donor. Conjugation frequencies for pAM771 under mouse intestine (B), planktonic (D), and biofilm (F) settings are also shown. Experiments with T11RF recipients are represented with squares; with T11RF Δ*cas9* recipients as triangles. For calculating *in vivo* conjugation frequencies, the conjugation frequency for each mouse was determined by dividing the transconjugant CFU/g by the donor CFU/g; one symbol represents one mouse on the graph. Black horizontal bars represent the geometric mean of data in each group. No symbol means that a frequency could not be calculated because one or both of the values (donor CFU/g or transconjugant CFU/g) were zero. Statistical significance was assessed using a two-tailed Student’s t-Test; *P*-values, *<0.05, **<0.01 and ***<0.001.

**Dataset S1.**
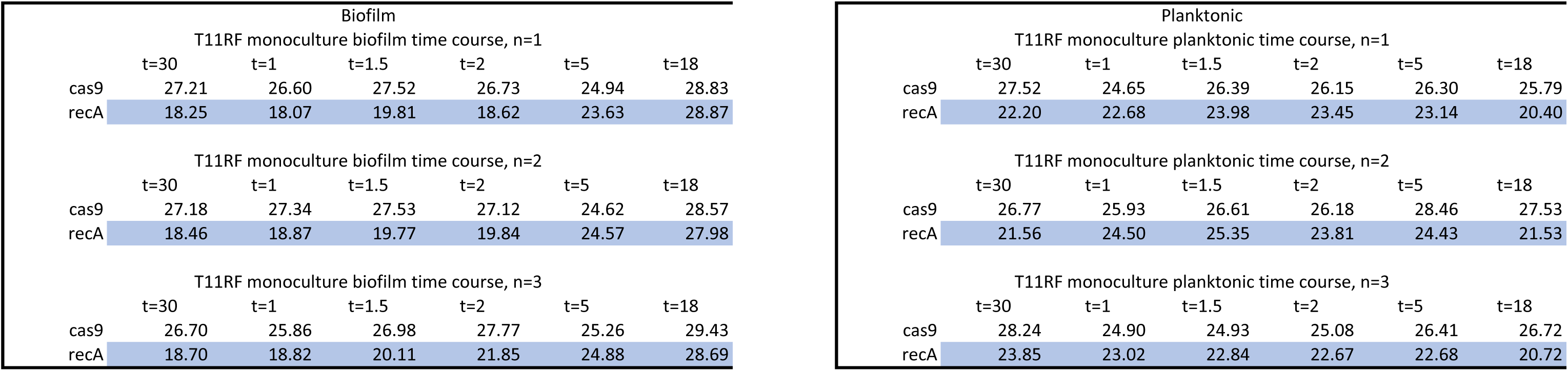
Ct values for RT-qPCR.

